# Risk Assessment of Arterial Embolism from Neofilera^®^ Filler in Rabbits

**DOI:** 10.64898/2026.06.08.730770

**Authors:** Vasanop Vachiramon, Tussapon Boonyarattanasoonthorn, Jeeraprapha Duangbupha, Anusak Kijtawornrat, Chia-Chang Liu, Chia-Hsien Hsieh

## Abstract

The incidence of vascular complications differs among dermal filler formulations. This study aimed to evaluate the embolic risk associated with Neofilera^®^, a filler composed of carboxymethyl cellulose and polylactic acid microspheres, following intra-arterial injection. The central auricular arteries of rabbits were injected with Neofilera^®^ at volumes of 0.1 mL or 0.2 mL under various conditions: normal saline (control) and Neofilera^®^ diluted at ratios of 1:5 (Group 1), 1:10 (Group 2), and 1:15 (Group 3). The presence of transparent emboli was assessed immediately after injection, while tissue necrosis (percentage and area) and histopathological alterations were evaluated on days 1 and 7 post-injection. Relative to controls, Neofilera^®^ administered at 0.1 mL dispersed within minutes and did not induce significant tissue necrosis at either observation time point. In contrast, administration of 0.2 mL, even in diluted form, was associated with an increased incidence of vascular occlusion. Overall, these findings indicate that Neofilera^®^ presents a lower embolic risk when injected at a volume of 0.1 mL, whereas higher injection volumes may substantially increase the likelihood of embolic complications.

## 1. Introduction

Minimally invasive cosmetic filler injections have become increasingly prevalent in aesthetic medicine due to their convenience and favorable cosmetic outcomes [1–4]; however, these procedures are not without risk [4–10]. Neofilera^®^ is a biodegradable tissue filler composed of carboxymethyl cellulose (CMC) combined with poly-D,L-lactic acid (PDLLA) microspheres, and is designed for both soft- and hard-tissue augmentation. CMC, a chemically modified cellulose derivative, exhibits high viscosity, excellent bio-compatibility, and favorable rheological properties, making it widely applicable in biomedical and reconstructive settings, including use as a filler material in implants and tissue scaffolds [11–14]. PDLLA, a synthetic biodegradable polymer, is well recognized for its capacity to stimulate collagen neogenesis and provide durable volumizing effects, with some formulations demonstrating clinical efficacy lasting up to 24 months [15–20]. The combination of CMC and PDLLA offers an effective aesthetic filler while meeting the growing demand for biodegradable, non-toxic, and bioresorbable materials in modern medical treatments [16,21,22].

Despite their widespread clinical use, injectable fillers carry inherent risks, particularly when inadvertent intravascular injection occurs [5,23,24]. Although the probability of such events is low, intravascular embolization may lead to severe and potentially irreversible complications [25–27]. Reported adverse outcomes following facial filler injections include visual disturbances, retinal artery occlusion, blindness, ischemic stroke, skin necrosis, temporary eschar formation, and permanent scarring [28–30]. Clinical manifestations may include acute changes in vision, neurological deficits consistent with cerebro-vascular events (such as facial drooping, limb weakness or numbness, speech impairment, dizziness, confusion, or severe headache), blanching or whitening of the skin, and disproportionate pain during or shortly after injection [5,31].

Despite these documented clinical complications, there remains a lack of controlled experimental studies evaluating the vascular risks associated specifically with Neofilera^®^. Therefore, the present study employs a rabbit model to investigate the hypothesis that intravascular exposure to filler materials may induce blood vessel occlusion and subsequent tissue ischemia or necrosis. The central auricular artery of each rabbit ear will be injected intra-arterially with Neofilera^®^ at defined volumes (0.1 or 0.2 mL) to simulate accidental intravascular administration during clinical filler injection. Key outcome measures will include filler dispersion patterns, changes in local vascular diameter, and the extent of tissue necrosis, providing critical insight into the vascular safety profile and potential risks associated with this filler material.

## 2. Materials and Methods

### 2.1. Animals

Forty healthy male and female white rabbits were maintained in standard rabbit cages with ad libitum access to food and water under a 12:12 h light–dark cycle in a temperature-controlled environment (20 ± 1 °C) with a relative humidity of 50 ± 10% at the Chulalongkorn University Laboratory Animal Center (CULAC, study number 2473032). All animals were humanely euthanized using carbon dioxide seven days following intra-arterial administration of the test material. The investigational product evaluated in this study was a dermal filler composed of CMC and PDLLA microspheres (Neofilera^®^; Diamond Biotechnology Co., Ltd., Taipei, Taiwan).

### 2.2. Experimental procedures

Four groups of rabbits were set up in this experiment: group control (normal saline), group 1 (1:5 dilution), group 2 (1:10 dilution), and group 3 (1:15 dilution). In group 1-3, the standard Neofilera^®^ solution was diluted with normal saline. Each group was subdivided into 2 subgroups: 0.1 mL/ear and 0.2 mL/ear, n=10/subgroup. The solution was injected via the central artery with either 0.1 mL or 0.2 mL of Neofilera^®^. The study design was summarized in Figure 1.

**Figure 1.**
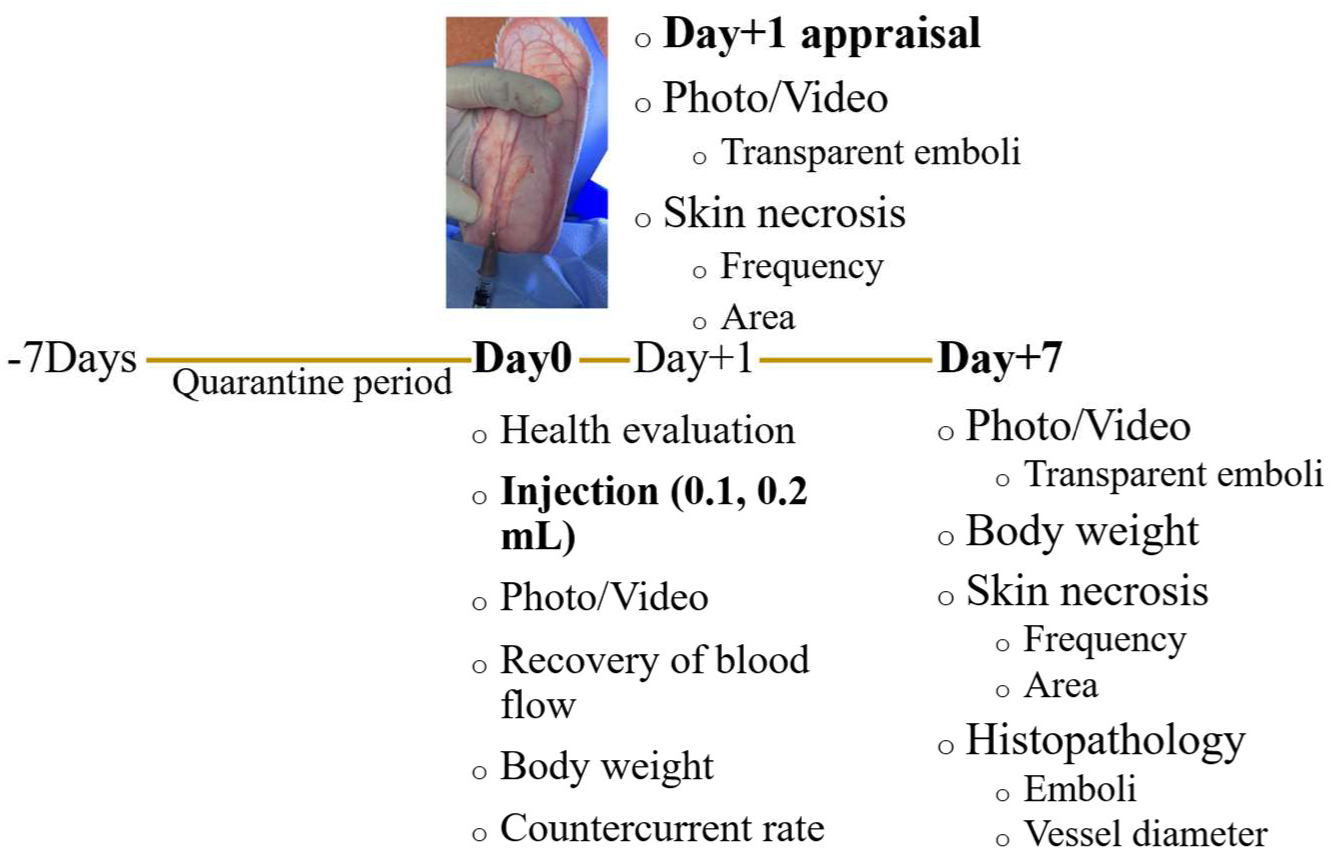
Schematic illustration of the study design.

On the day of the experiment, rabbits were gently restrained using a standard rabbit restrainer. The intended injection site on each ear was shaved and initially treated with topical lidocaine spray; the area was subsequently cleansed with saline-soaked gauze 15–20 minutes later. The ventral surface of the ear was transilluminated using a high-intensity light source to visualize the vascular structures. The test filler was then administered in-tra-arterially into the main trunk of the central ear artery (CEA) using a 26-gauge needle. The injection point was located approximately 1 cm proximal to the medial branch of the central ear vein (CEV), corresponding to about 4–5 cm from the base of the ear (Figure 2). The injection rate was carefully controlled at approximately 0.2 mL/min.

**Figure 2.**
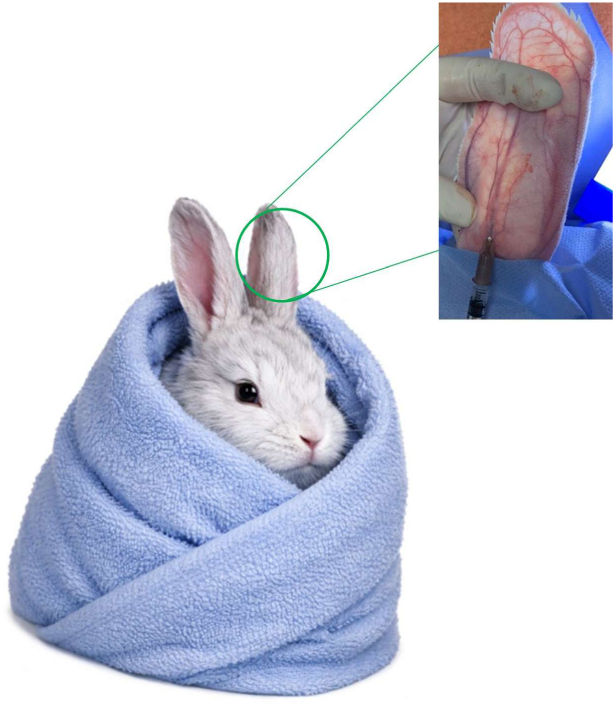
Schematic representation of the principal vasculature of the rabbit ear, indicating the designated injection site, which is located approximately 1 cm proximal to the medial branch of the central ear vein.

### 2.3. Observation of CEA Blood Flow and Skin Necrosis

Blood flow was visually assessed and documented using a digital camera prior to injection and within 5 minutes following administration. The 5-minute post-injection time point was selected to evaluate early vascular recovery based on previously published studies [32]. The countercurrent rate was defined as the proportion of ears exhibiting retrograde flow of the injected filler beyond the injection site. In addition, vascular perfusion, skin coloration, and the extent and area of tissue necrosis were systematically evaluated and photographed on days 1 and 7 post-injection for each treated ear.

### 2.4. Histopathological Changes of Blood Vessels on Day 7 Post-Injection

The entire ear pinna was harvested and fixed in 10% neutral-buffered formalin. Following adequate fixation, each ear was sectioned into three regions including proximal, middle, and distal along the course of the central ear artery (Figure 3). The tissue samples were subsequently processed using standard histological procedures, embedded in paraffin, sectioned at a thickness of 4 µm, and stained with hematoxylin and eosin (H&E) for microscopic evaluation.

**Figure 3.**
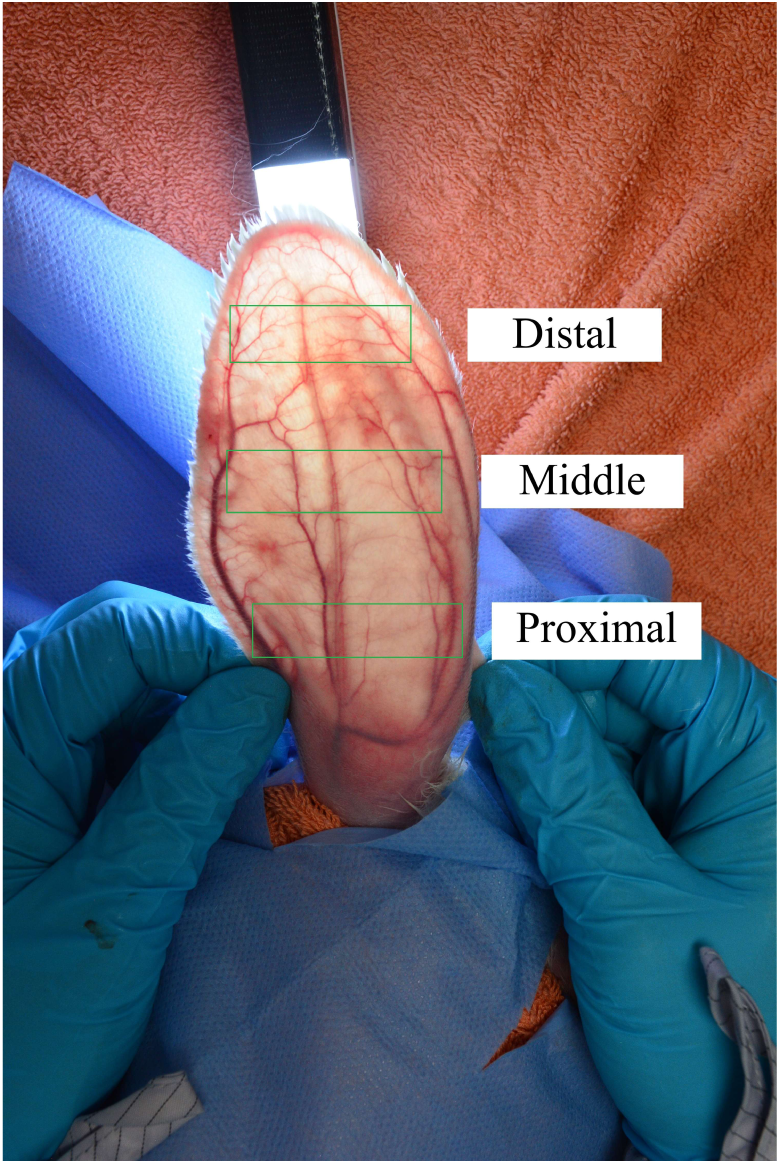
Schematic representation of tissue sectioning of the rabbit ear. The green dashed boxes indicate the locations of the proximal, middle, and distal segments, respectively.

The histopathological changes were observed using light microscope (Nikon Eclipse Ci microscope; Nikon, Tokyo, Japan) and photographed with microscope equipped Nikon DS-Ri2 camera (Nikon). The diameter of the largest vessel in each distal tissue section was measured as previously described [32]. Image analysis was performed using NIS-Elements BR software version 5.21.00 (Nikon).

### 2.5. Statistical Analysis

All data are analyzed and expressed as mean ± standard deviation. Chi-square test was used to evaluate frequency and percentage. A nonparametric test was used to analyze the area of skin lesion and diameter of the largest vessel. Two-way ANOVA was used to calculate body weight. P-values < 0.01 were considered significant.

## 3. Results

### 3.1. Immediate Effects of Intra-Arterial Injection of Neofilera^®^ on Blood Vessels

No statistically significant differences in body weight were observed among the eight groups. The percentage change in body weight from baseline to 7 days after Neofilera^®^ injection is shown in Figure 4, indicating that all rabbits exhibited weight gain over the course of the study.

**Figure 4.**
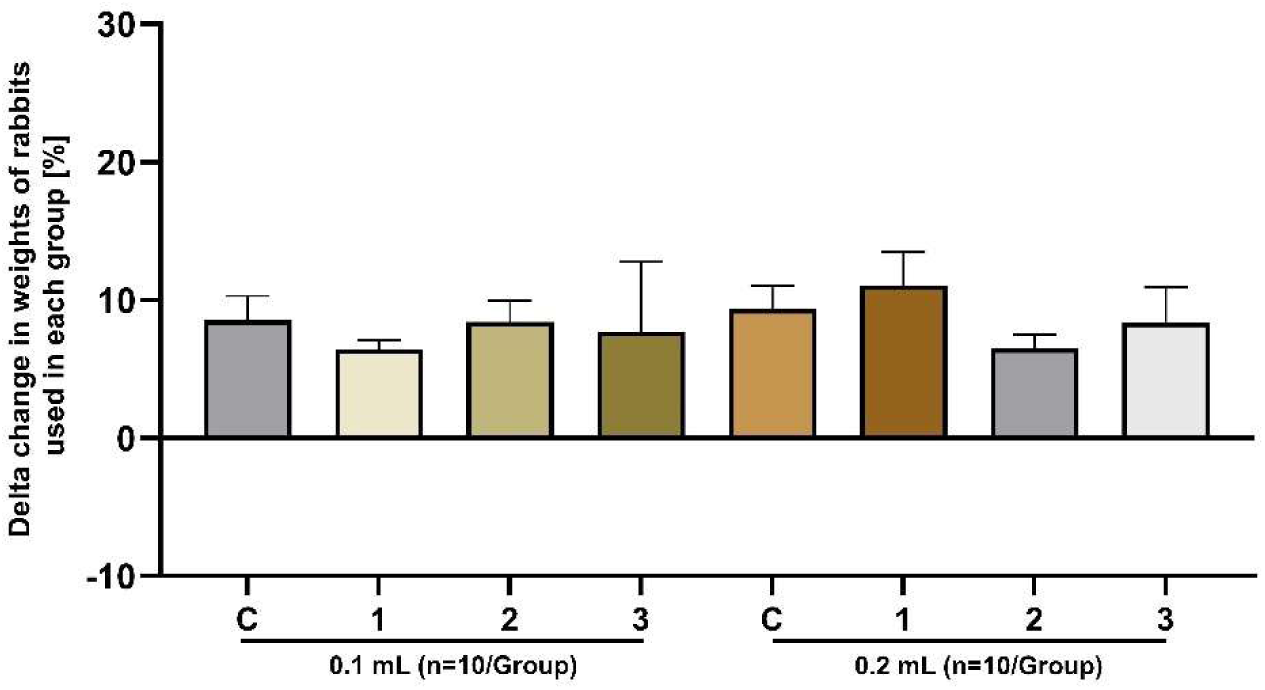
Percent change in body weight of rabbits used in each group (n=10/group).

Before injection, the vascular structures of the rabbit ear were readily visualized under strong transillumination (Figure 5). The distal branches of the central ear artery displayed a characteristic Y-shaped pattern, allowing clear identification of both the main arterial trunk and the accompanying central vein. During administration, the injected filler progressively spread within the arterial system, extending along the primary trunk as well as the collateral branches of the central ear artery. The incidence of different blood flow responses immediately following intra-arterial injection of Neofilera^®^ is summarized in Figure 6. Blood flow responses were classified into three categories: Category I, indicating complete dispersion of the filler without vascular occlusion; Category II, representing transient arterial occlusion with near-complete recovery of blood flow within 5 minutes; and Category III, indicating persistent arterial occlusion with incomplete restoration of blood flow within 5 minutes post-injection.

**Figure 5.**
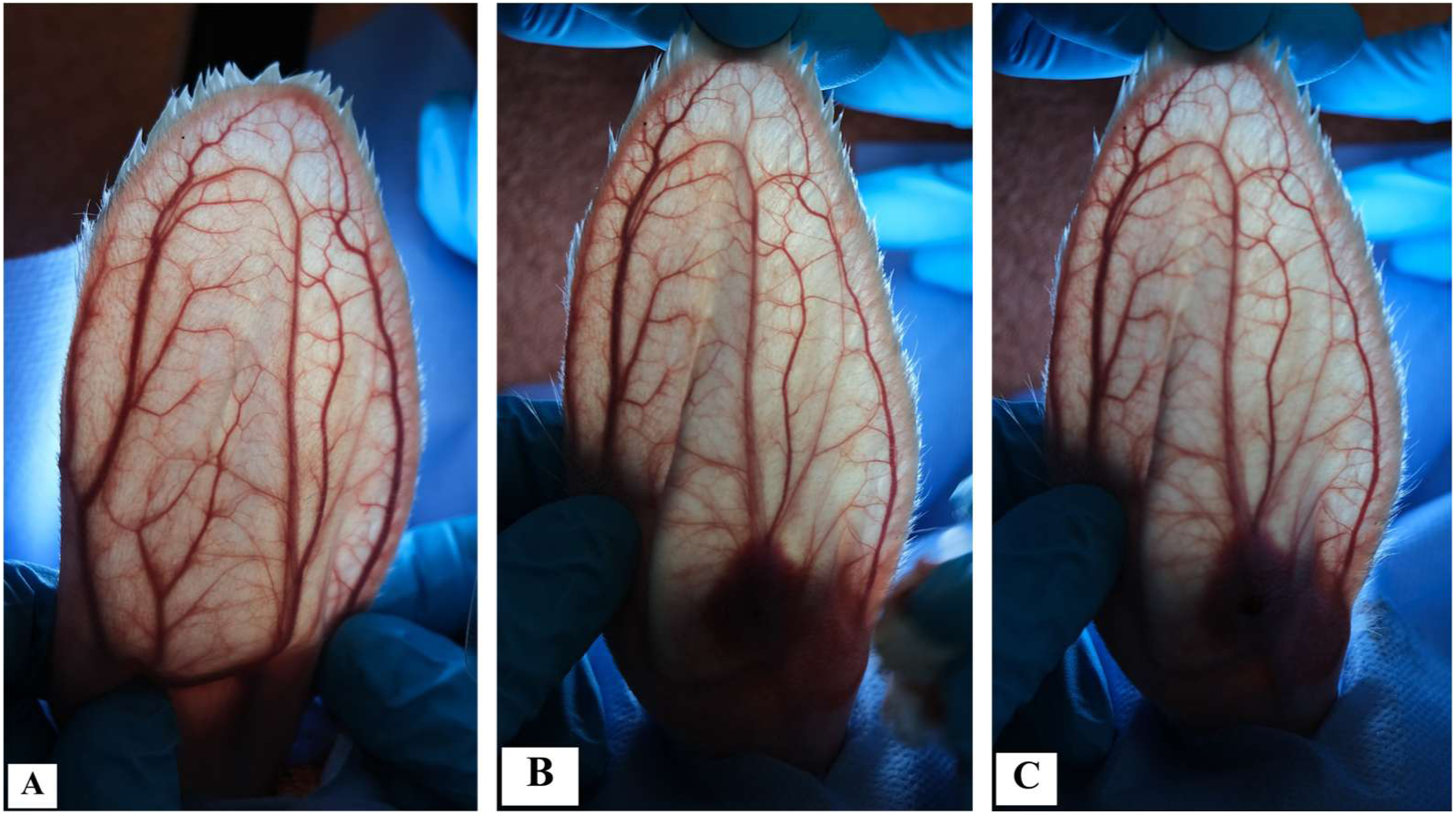
Characteristics of blood vessels of the rabbit ear before and after injection of 0.2 mL Neofil-era^®^. A) before injection, B) right after injection, and C) 5 minutes after injection.

**Figure 6.**
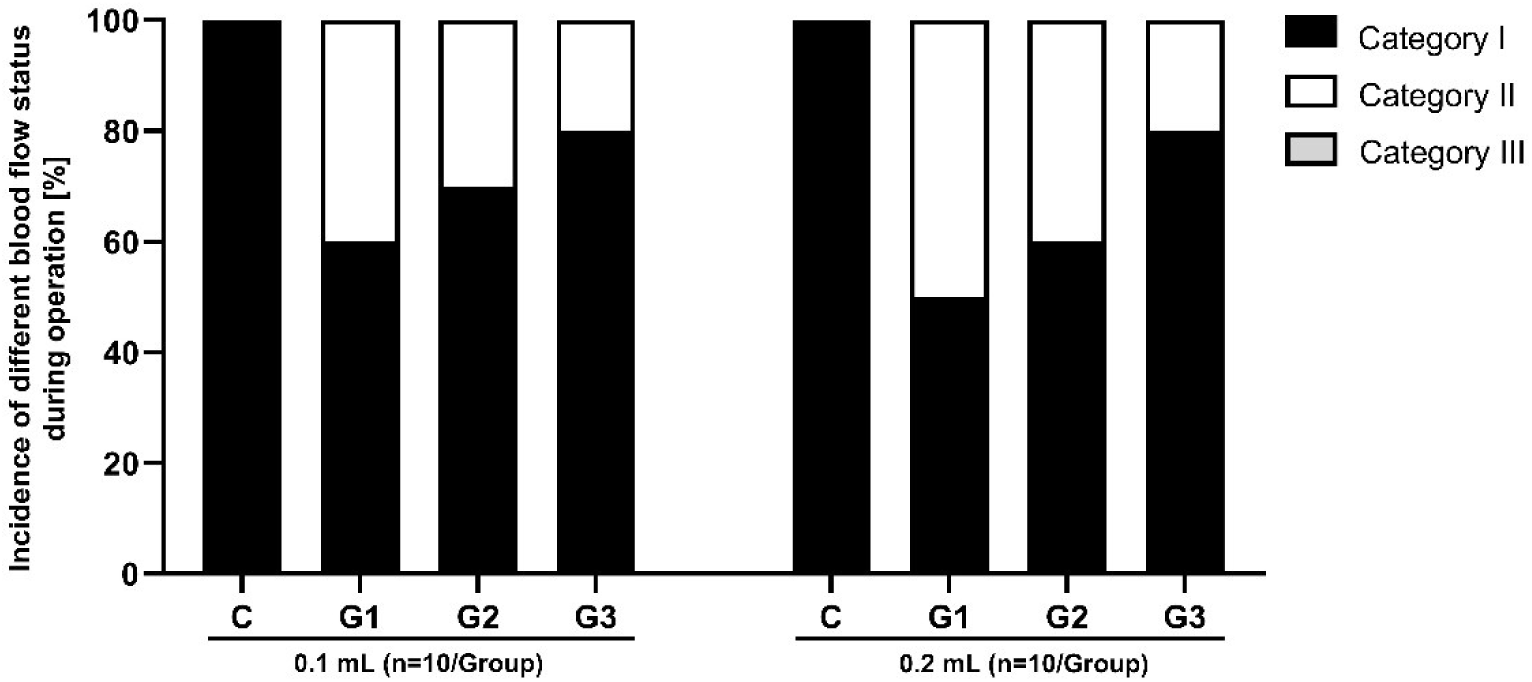
Incidence of blood flow status within 5 minutes of Neofilera^®^ injection. Black = category I (no block), white = category II (temporarily block but restored blood flow within 5 minutes, grey = category III (sustained blockage of the main branch of central ear artery beyond 5 minutes after injection) *P < 0.05 when compared with its control group, ^#^P < 0.05 when compared between groups (i.e., 0.1- vs 0.2-mL) at the same group, N=10/group

For the 0.1 mL injection volume, the control group exhibited 100% Category I responses, indicating normal blood flow without evidence of embolism. In the diluted Neofilera^®^ groups, Category I responses remained predominant, although a gradual decrease in Category II responses was observed with more dilution ratios. Notably, no Category III responses were observed across all dilution groups at the 0.1 mL injection volume, suggesting a low risk of sustained vascular occlusion at this dose. In contrast, administration of Neofilera^®^ at a volume of 0.2 mL resulted in a higher incidence of adverse blood flow responses. While the control group again demonstrated 100% Category I responses, Group 1 (1:5 dilution) and Group 2 (1:10 dilution) showed a marked increase in Category II responses. The incidence of Category II responses was notably higher in Group 1 and group 2 of the 0.2 mL injection compared with the corresponding 0.1 mL group, indicating a volume-dependent increase in embolic risk.

The countercurrent rate observed during intra-arterial injection is summarized in Figure 7. In the 0.1 and 0.2 mL injection group, the control group exhibited no countercurrent rate, indicating no reflux of the injected material. In contrast, Groups 1 and 2 showed a noticeable increase in countercurrent events, each demonstrating an incidence of approximately 30 – 40% comparable to the control group. Increasing dilution was associated with a reduction in countercurrent incidence, indicating that both injection volume and filler concentration influence the likelihood of reflux during intra-arterial administration.

**Figure 7.**
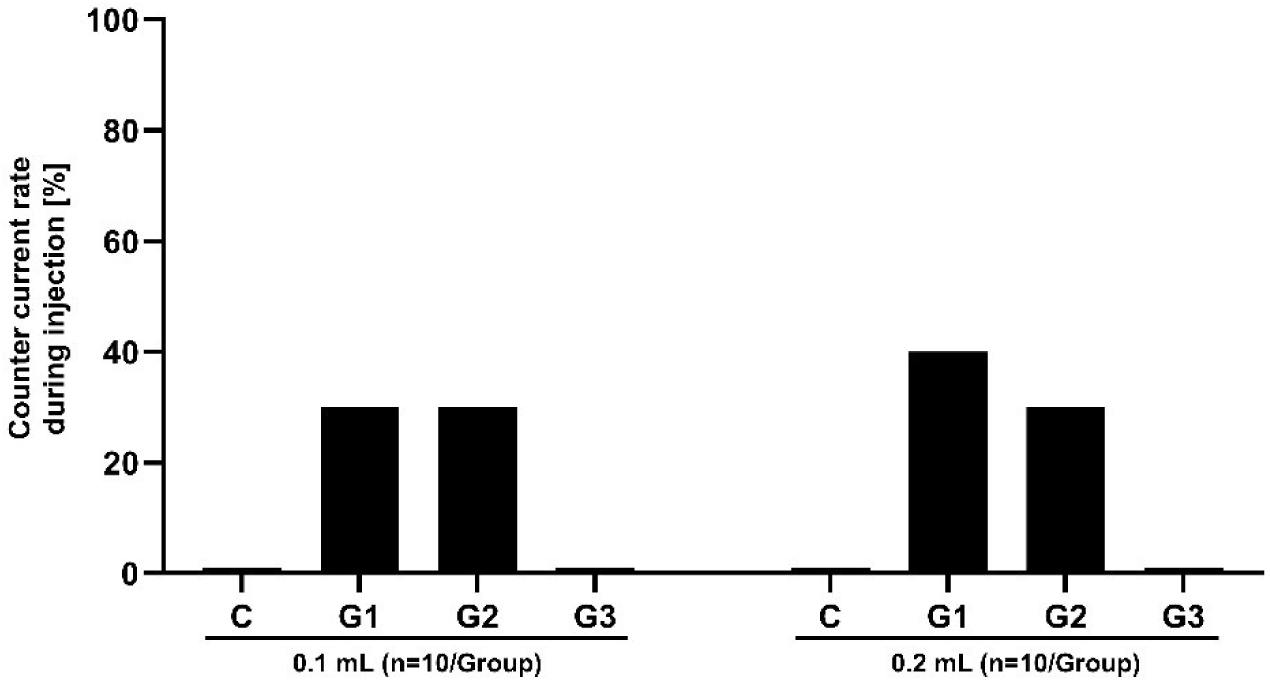
Countercurrent rate during injection. *P < 0.05 when compared with its control group, ^#^P < 0.05 when compared between groups (i.e., 0.1- vs 0.2-mL) at the same group, N=10/group

This section may be divided by subheadings. It should provide a concise and precise description of the experimental results, their interpretation, as well as the experimental conclusions that can be drawn.

### 3.2. Short-Term Effects of Intra-Arterial Injection of Neofilera^®^ on Vascular Occlusion and Skin Damage

On day 1 after injection, no transparent emboli, tissue necrosis, or measurable necrotic areas were detected in any of the injected rabbit ears, including both the control and Neofilera-treated groups (Figure 8). Accordingly, no differences were observed between rabbits receiving 0.1 mL and those receiving 0.2 mL of the filler. Furthermore, varying the dilution of Neofilera^®^ did not influence the occurrence of emboli or the extent of tissue necrosis on day 1 post-injection (Figures 9–11). On day 7 post-injection, the vascular occlusion status was similar to that of day one post-injection (i.e., no occlusion in all rabbit groups).

**Figure 8.**
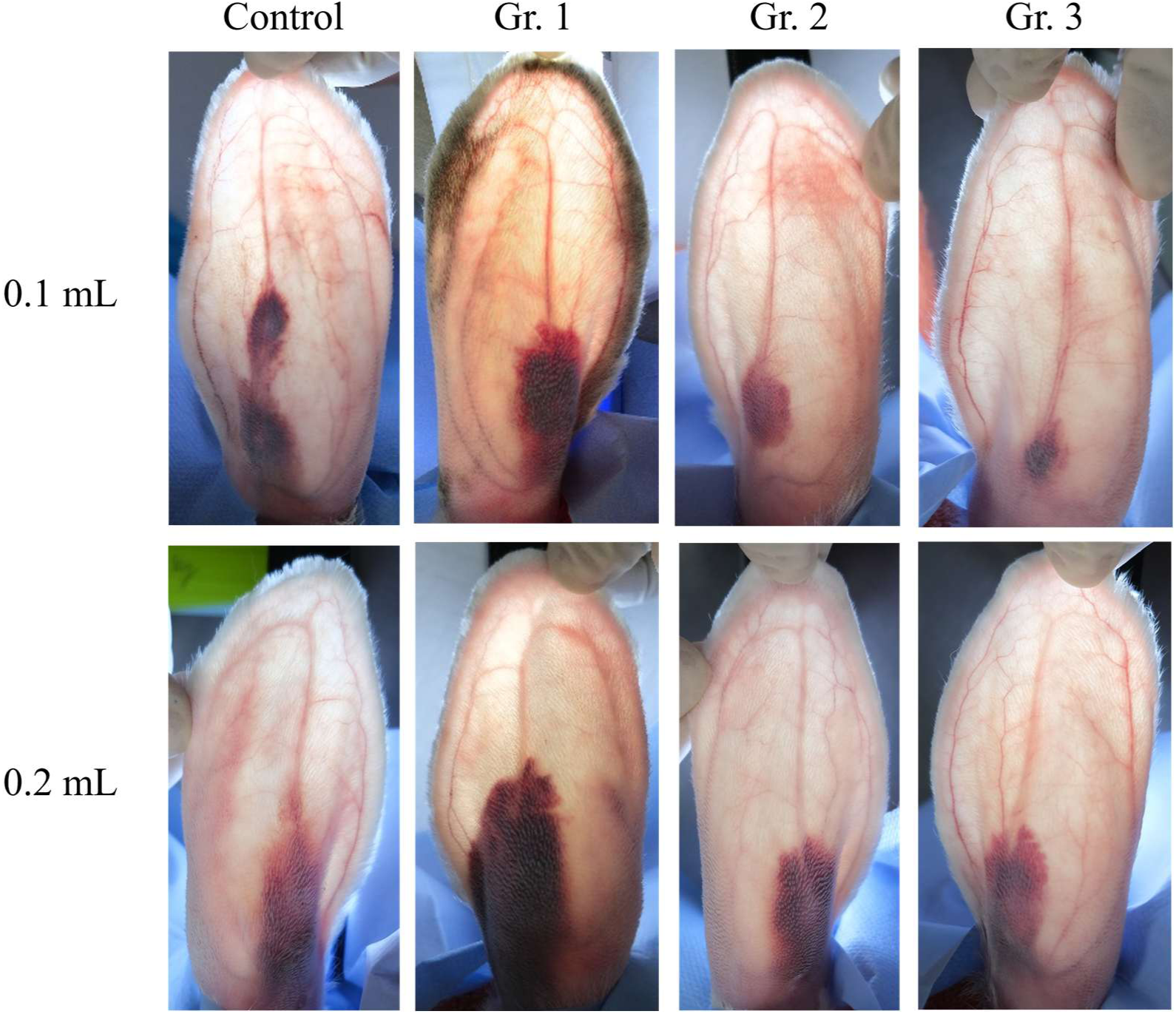
Representative photo of rabbit’s ear observed on day 1 post-injection.

**Figure 9.**
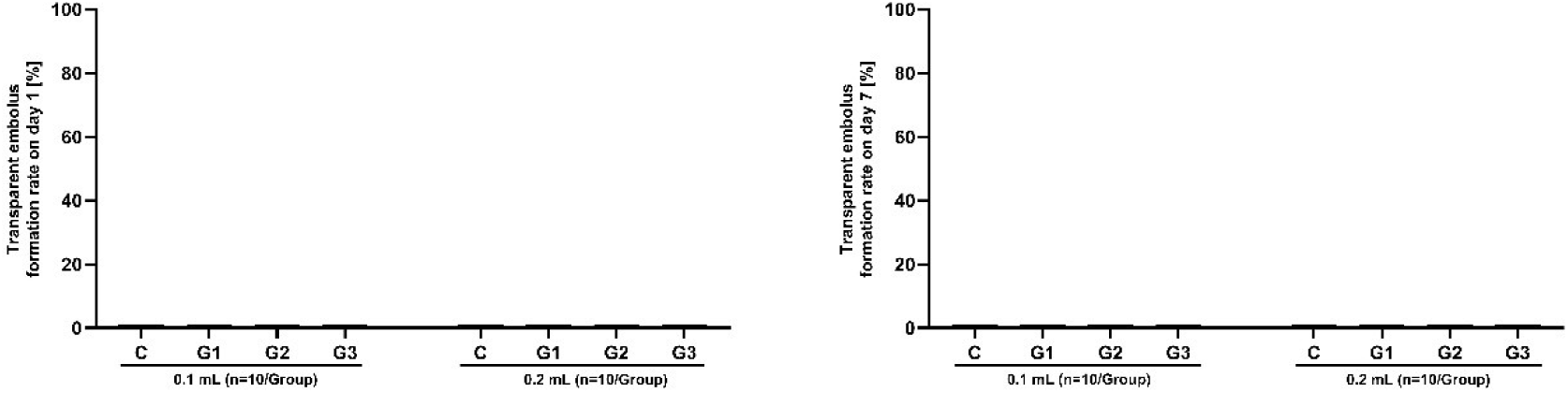
Transparent embolus rate on day 1 and day 7 post-injected Neofilera^®^ (n=10/group).

**Figure 10.**
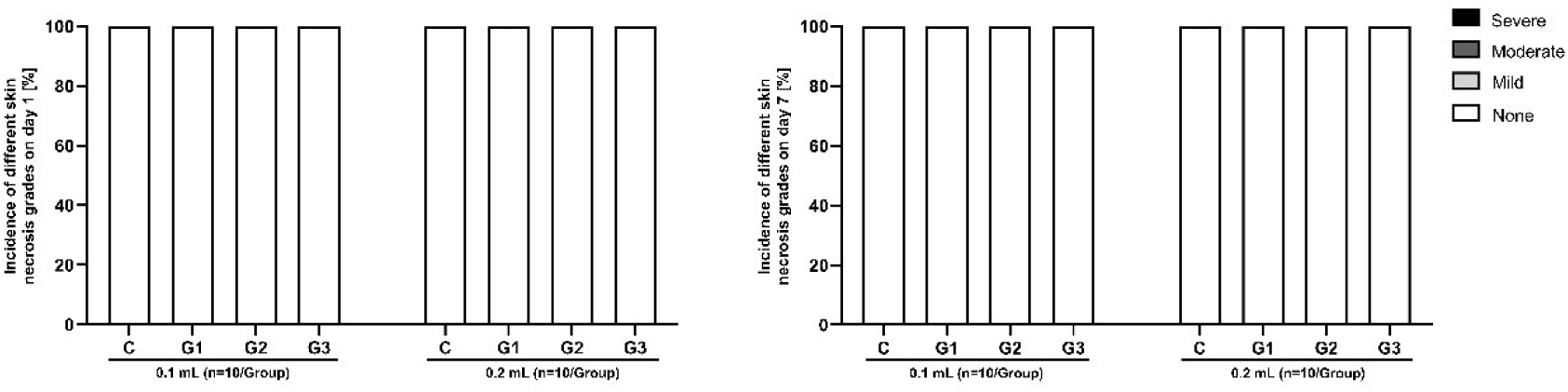
Skin necrosis grade on day 1 and day 7 post-injected Neofilera^®^ (n=10/group).

**Figure 11.**
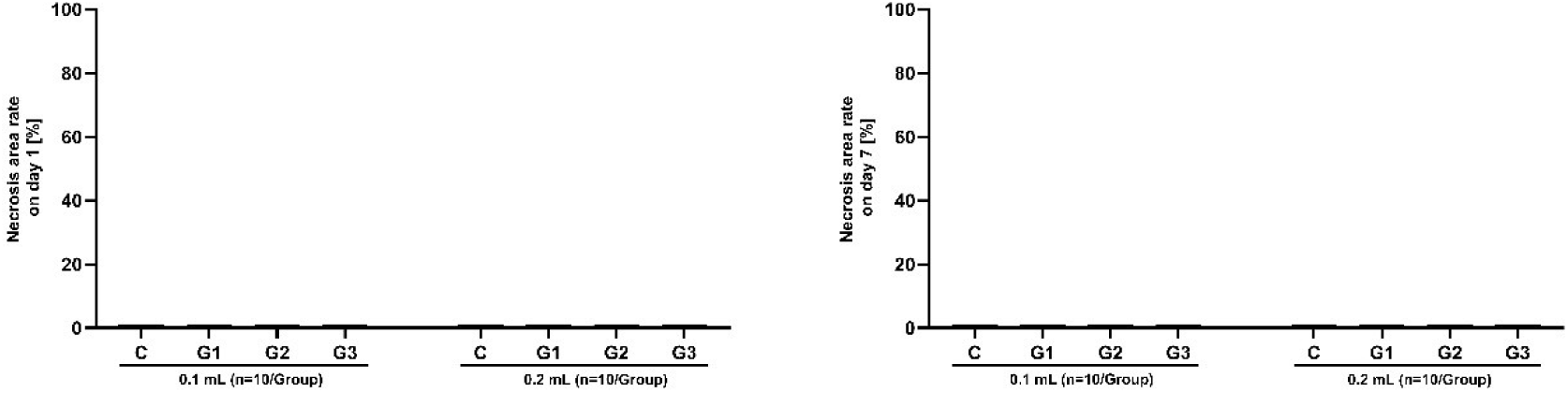
Necrosis area rate on day 1 and day 7 post-injected Neofilera^®^ (n=10/group).

### 3.3. Histopathological Changes of Blood Vessels on Day 7 Post-Injection

Mostly proximal segment sections of every group of both 0.1- and 0.2-mL injection demonstrated no thrombi or emboli found in all vessels in both left and right ear sections. There are necrotic foci and inflammatory cells infiltration (predominantly macrophages and lymphocytes) adjacent to the vessels of both left and right ear sections (Figures. 12-14). Hemorrhage is also found in one section of left ear.

**Figure 12.**
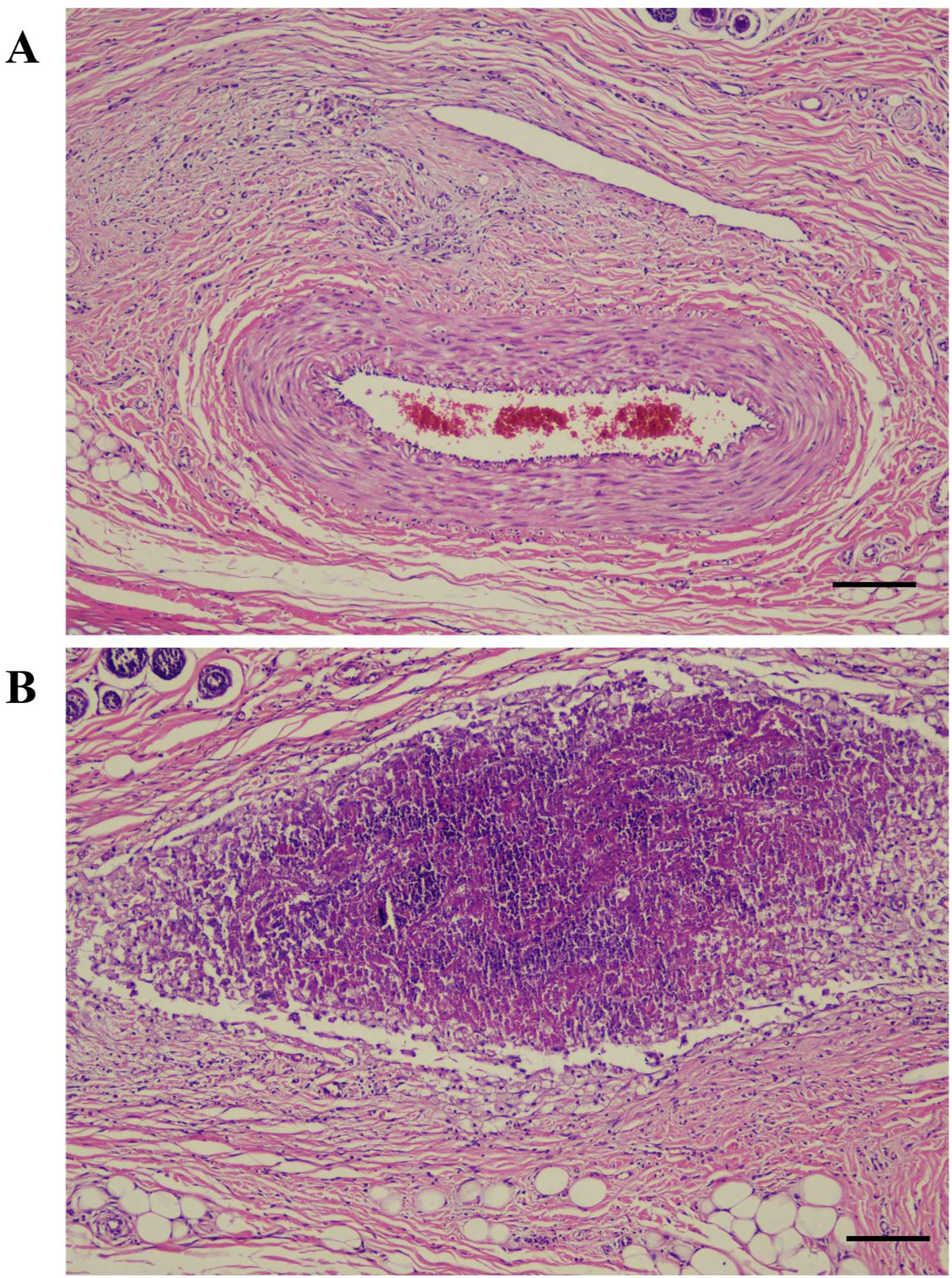
Proximal segment of the ear vessel in Group 1 (1:5 dilution) following injection of Neofil-era^®^ diluted with normal saline solution (NSS). (A) Inflammatory cell infiltration and (B) necrotic area in the perivascular tissue. Scale bar = 100 µm.

**Figure 13.**
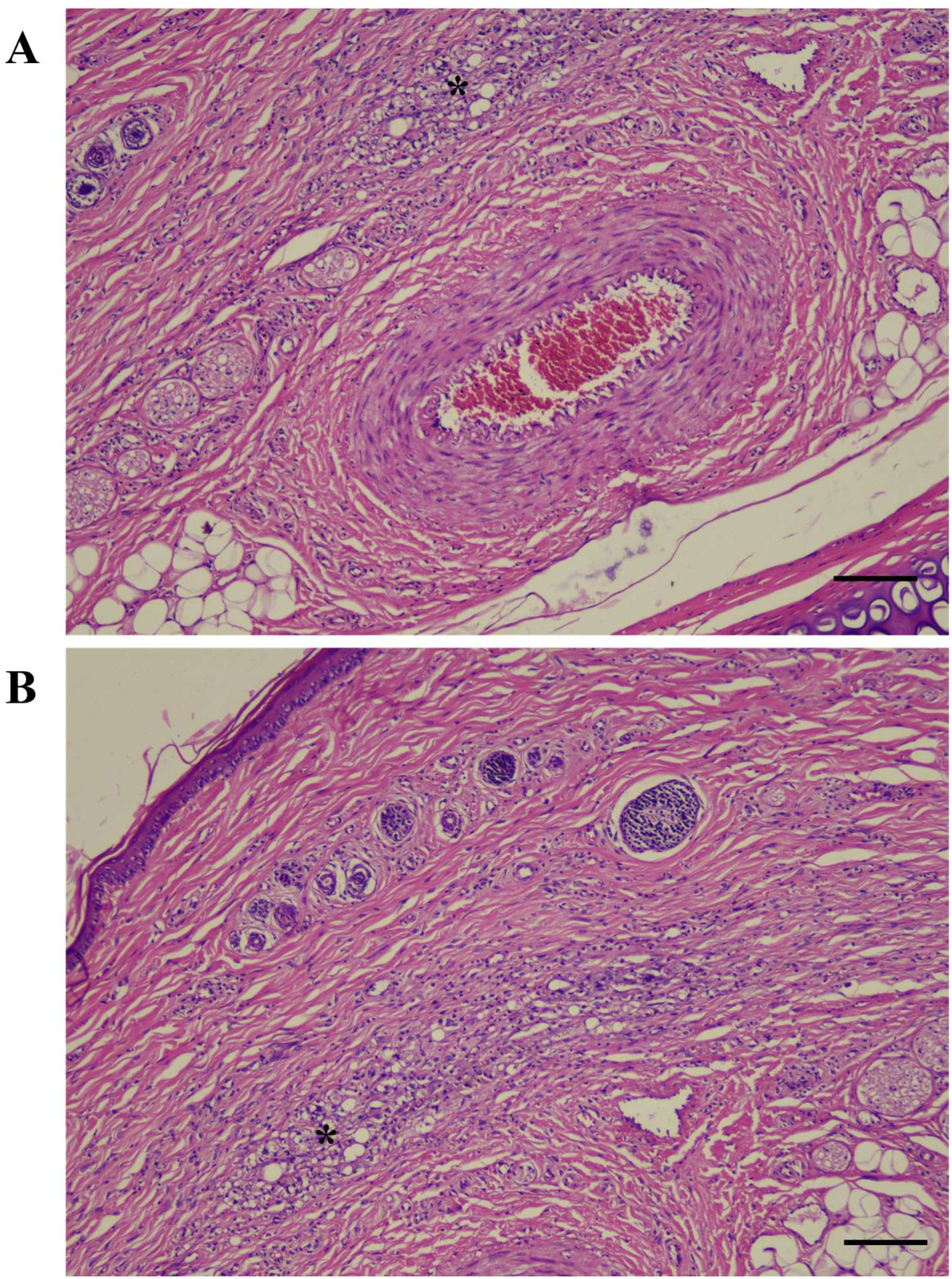
Proximal segment of the ear vessel in Group 2 (1:10 dilution) following injection of Neofil-era^®^ diluted with normal saline solution (NSS). (A and B) Inflammatory infiltration in the adjacent area of vessel (*). Scale bar = 100 µm.

**Figure 14.**
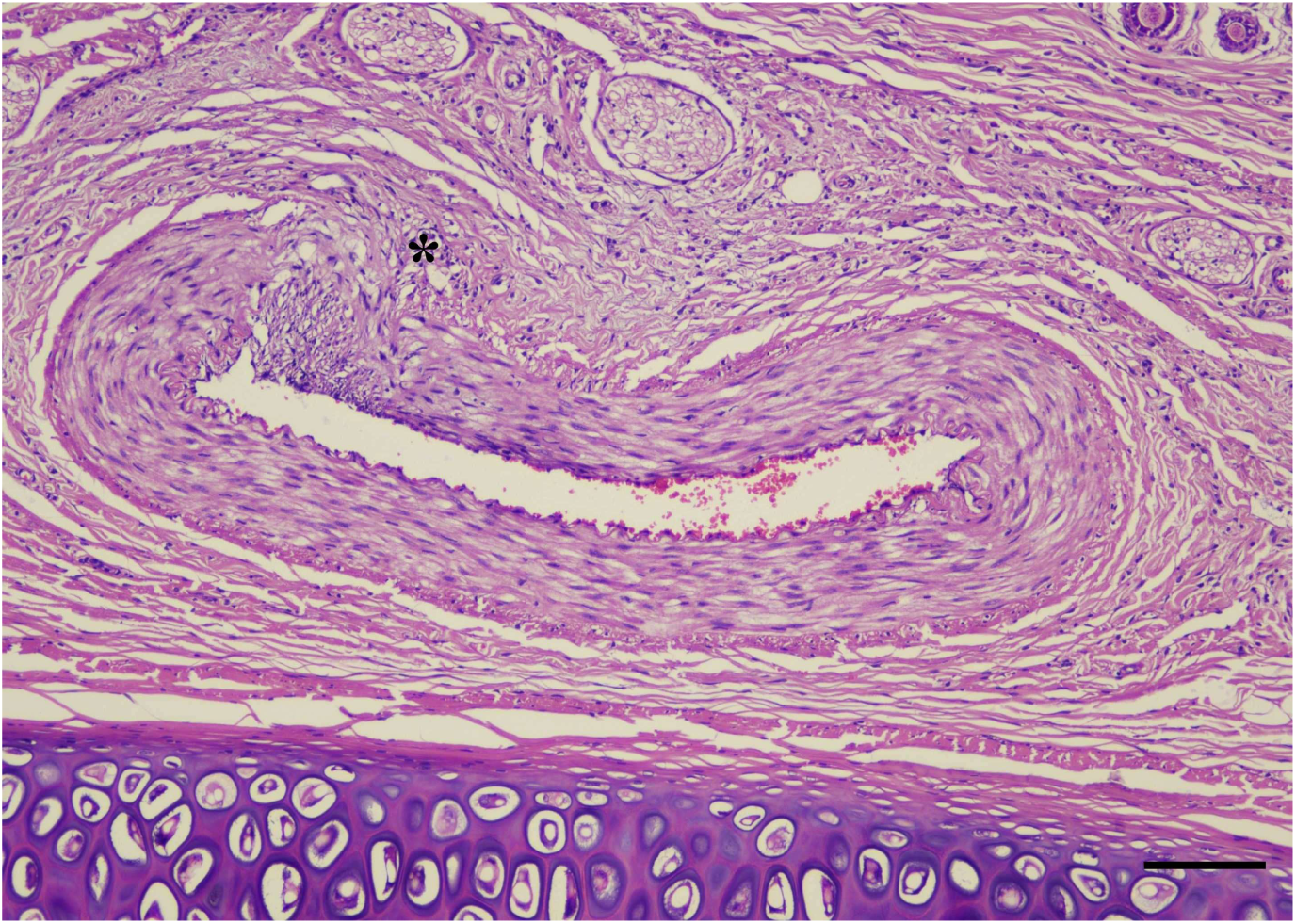
Proximal segment of the ear vessel in Group 3 (1:15 dilution) following injection of Neofil-era^®^ diluted with normal saline solution (NSS). Inflammatory infiltration in the adjacent area of vessel (*). Scale bar = 100 µm.

The middle segment sections (both 0.1 ml and 0.2 ml injection) demonstrated no thrombi or emboli found in all vessels in both left and right ear sections. For the distal segment, there was no inflammation, necrosis, hemorrhage, thrombosis and embolization were found in every group including the control group. The representative of histological sections of every group are showed in Figure 15 and the diameters of the largest vessels in distal segments are showed in Table 1.

**Figure 15.**
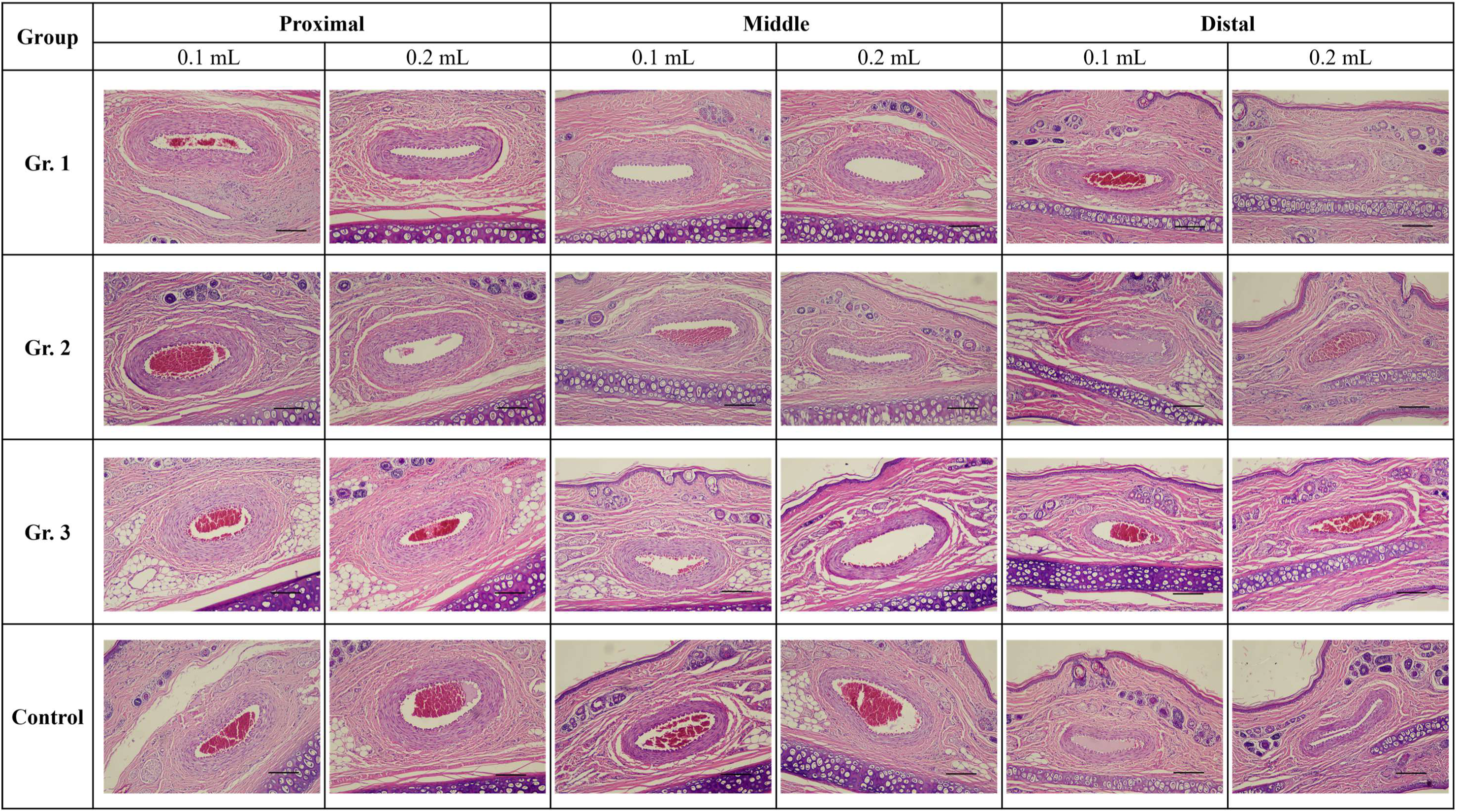
Representative vessels located at the proximal, middle, and distal segments of the rabbit ear after injection of Neofilera^®^. Scale bar = 200 µm.

**Table 1.**
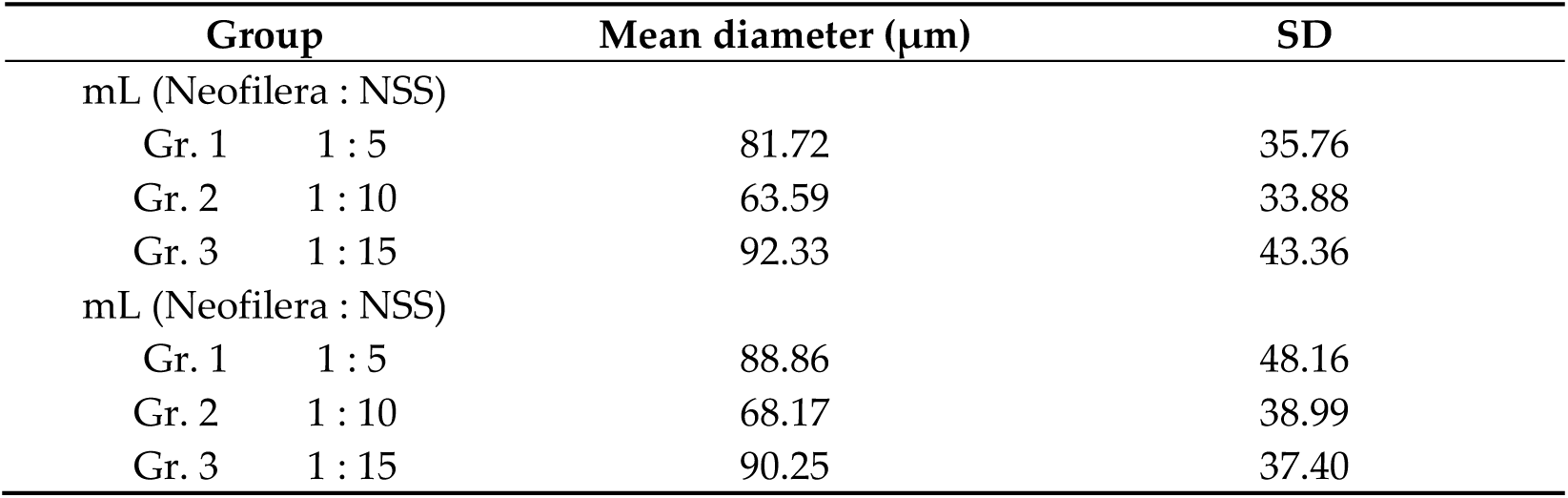
Diameters of the largest vessels in distal segment (n=10/group).

## 4. Discussion

This study investigated the risk of vascular embolism and tissue necrosis following inadvertent intra-arterial injection of Neofilera^®^, a biodegradable dermal filler composed of CMC and PDLLA, using a rabbit ear model. To our knowledge, this is the first controlled in vivo study to evaluate the embolic potential of Neofilera^®^ and the influence of injection volume and dilution on vascular safety. The results indicate that Neofilera^®^ injected at a volume of 0.1 mL posed minimal embolic risk, whereas increasing the injection volume to 0.2 mL increased the incidence of transient vascular flow disturbances, demonstrating a volume-dependent effect.

The rabbit ear model was chosen due to its thin skin and superficial vascular network, which allow direct visualization of embolic events and blood flow changes following filler injection [32]. Vascular occlusion is recognized as one of the most severe complications associated with cosmetic fillers, potentially leading to ischemia, tissue necrosis, and permanent functional impairment if not promptly managed [7,23,28,33]. In this study, immediate post-injection observations showed that 0.1 mL injections, regardless of dilution, resulted primarily in complete filler dispersion without arterial occlusion, while higher-vol-ume injections were associated with increased transient arterial blockage.

Importantly, increasing the dilution of Neofilera^®^ was associated with a reduction in countercurrent flow and embolic incidence, suggesting that lower filler viscosity and particle concentration may facilitate dispersion and reduce the likelihood of arterial obstruction. This observation is consistent with previous studies demonstrating that dilution of particulate fillers can mitigate embolic risk by decreasing particle aggregation and improving intravascular flow dynamics [32]. The physicochemical properties of CMC and PDLLA both biodegradable and biocompatible polymers may further contribute to the relatively low persistence of vascular occlusion observed in this study [12,16].

Despite the transient vascular disturbances observed immediately after injection, no transparent emboli, macroscopic skin necrosis, or significant necrotic areas were detected on days 1 and 7 post-injection across all groups. Histopathological analysis revealed focal perivascular necrosis and inflammatory cell infiltration, predominantly macrophages and lymphocytes, primarily confined to proximal segments of the ear vasculature. These findings likely reflect localized tissue responses to mechanical vascular insult or temporary ischemia rather than sustained embolic obstruction. Notably, no thrombi or emboli were identified in any vascular segment, and distal segments showed no evidence of inflammation or necrosis, further supporting the conclusion that Neofilera^®^ does not induce persistent arterial occlusion under the conditions tested.

From a clinical perspective, these results underscore the importance of injection technique, volume control, and filler preparation in minimizing vascular complications. Although Neofilera^®^ demonstrated a favorable safety profile at lower injection volumes, inadvertent intravascular injection of higher volumes may still pose a risk of transient embolic events. Therefore, adherence to safe injection practices, thorough anatomical knowledge, and consideration of dilution strategies remain essential for reducing adverse outcomes [24,34].

In conclusion, Neofilera^®^ demonstrates a favorable vascular safety profile when injected at lower volumes, with increased embolic risk observed at higher injection volumes. Dilution of the filler appears to further reduce this risk. These findings highlight the importance of volume control, appropriate dilution, and meticulous injection technique to minimize vascular complications associated with dermal fillers.

## Funding

This research was funded by Diamond Biotechnology Co., Ltd., grant number 2026-N-006.

## Informed Consent Statement

Not applicable.

## Data Availability Statement

The data presented in this study are available upon request from the corresponding author.

## Conflicts of Interest

This work was supported by Diamond Biotechnology Co., Ltd. The corresponding author Chia-Hsien Hsieh is the Chairman and a shareholder of Diamond Biotechnology Co., Ltd. The other corresponding author Chia-Chang Liu is the CTO of Diamond Biotechnology Co., Ltd. The collection, analysis, independent interpretation of data, and manuscript writing were primarily conducted by the other co-authors to ensure academic objectivity. All authors declare no other competing interests.

## Abbreviations

The following abbreviations are used in this manuscript:

CMC: carboxymethyl cellulose
PDLLA: poly-D,L-lactic acid
CEA: central ear artery
CEV: central ear vein
H&E: hematoxylin and eosin
NSS: normal saline solution<colcnt=1>

